# Torque-optimized body morphing enables rapid surface to underwater diving in whirligig beetles

**DOI:** 10.64898/2026.09.29.755344

**Authors:** Yukun Sun, Emily Palmer, Lirong Xiang, Jena Shields, Aaron Quan, Sunghwan Jung, Chris Roh

## Abstract

Known for their rapid water surface locomotion, whirligig beetles (*Gyrinidae*) are also fast divers. They can transition from the surface to underwater swimming in under 100 ms. Diving requires them to generate sufficient hydrodynamic torque to break the water surface and reorient the body. However, the mechanism by which whirligigs create the necessary pitch-down torque is unknown. Here, we show that their diving is achieved by bending the abdomen. The inertial hydrodynamic pressure on the flexed abdomen generates the torque necessary for surface breaking and body rotation. The torque is maximized at an abdominal angle of 45°, whereas for effective diving, whirligigs display a preferential abdominal angle of around 28°, closely aligning with the angle that maximizes the torque-to-drag ratio. This abdominal angle is optimized by balancing torque generation with the drag penalty. Our findings inspire a mechanism for surface-to-underwater transition, offering design principles for multimodal robots capable of locomotion across the air-water interface.

## INTRODUCTION

The whirligig beetle (*Gyrinidae*), the fastest swimming insect on the water surface (*1*–*5*), rapidly disappears underwater when aerial threat looms. They are highly effective divers, transitioning into full submergence in less than 100 ms (*5*–*11*). While prior biomechanical studies on whirligigs have focused predominantly on the hydrodynamics of their horizontal surface swimming, the mechanics of their diving transition remains poorly understood. A diving transition from an aerial or semi-aquatic environment to underwater is important for multimodal animals and robots alike (*12*–*16*). Such a transition allows animals to explore a new fluid domain, providing resources such as prey capture (*14, 15, 17*), oviposition (*18*) and safety (*5*–*11*). Crucial to the whirligig’s survival, diving provides an effective escape strategy, alternative to high-speed surface swimming, enabling the beetle to anchor beneath the surface and remain concealed without continued energy-expenditure of active locomotion.

Typically, diving in biology requires generation of sufficiently high vertical momentum to break through and travel away from the air-water interface. The most common examples are plunge-diving birds, which readily break through the surface by dropping vertically before entering the water (*14, 15, 17*). However, unlike plunge divers, surface-dwelling whirligigs cannot vertically accelerate using gravity. Furthermore, the very adaptations that allow them to remain afloat on the water surface (*e*.*g*., buoyant body, pinned surface tension and flotation hairs) become sources of considerable resistance to their diving maneuver. Despite these limitations, the whirligigs dive successfully by breaking the surface and reorienting their bodies. However, the origin of the torque (*i*.*e*., pitching moment) that is necessary for both penetrating through the water surface and reorienting the body is still unclear.

Their diving as a startle response has been documented more than a century ago by Miall in 1895 (*6*). But the mechanism of their dive was first offered by Brown and Hatch in 1929 (*9*), highlighting that this transition required turning the body about a “horizontal transverse axis”, i.e., a pitching maneuver. They presumed that a change in leg kinematics would bring about the necessary maneuver for diving. The use of appendages for diving resurfaced in recent work, which suggested that the whirligig beetle relies on leg-based thrust vectoring (*5*), a mechanism observed in other similarly sized aquatic insects (*19, 20*). The study, however, does not show how the leg kinematics change to generate the required torque, leaving this mechanism to be further verified.

Alternatively, the torque could arise from a morphing body. If a body conformation change is the source of the torque, it is likely that the torque is generated by the wetted surface protruding into the water, as interacting with water (approximately 800 times denser than air) would provide a significantly greater fluid dynamic force. Earlier observations proposed anterior body morphing, suggesting that the beetle bends its head and prothorax downward to initiate the pitch (*3*). However, it is unclear what methodology was used to observe the body conformation changes during the rapid diving. Furthermore, it is uncertain whether anterior flexing truly occurs, and if it sufficiently protrudes the head below the water surface to generate the necessary torque. Otherwise, it is possible that different morphable body parts might be in use.

The whirligig beetle’s elusive diving mechanism remains a longstanding mystery, since robust observation of their rapid kinematics requires modern tools such as high-speed cameras. High-frame-rate recording has revealed much about their free surface swimming, as it occurs regularly even in tethered organisms. In contrast, capturing the unpredictable diving process requires identifying the exact moment the beetle submerges. This demands extended, continuous high-frame-rate recording capabilities that were unavailable at the time many of the earlier studies were conducted.

Here, using high-speed videography to capture these rapid transitions, we show that diving whirligigs generate torque for surface penetration and reorientation by morphing their abdomen (posterior section of the body). Flow visualization demonstrates that this posterior bending of the wetted surface redirects the oncoming flow, generating the hydrodynamic torque needed to pitch the body, pushing the head through the surface. Interestingly, while maximum torque is generated at a 45° abdominal angle, whirligigs display a preferential abdominal angle of approximately 28°. Our mechanistic models reveal that this preferred angle coincides with the angle that maximizes the torque-to-drag ratio. This optimization delicately balances the trade-off between generating the necessary pitching torque from lift force and minimizing the hydrodynamic drag penalty. These findings are expected to be translated into multimodal robotic systems.

## RESULTS

### Overall Diving Kinematics

The whirligigs’ diving kinematics (Fig. 1; Movie S1) were filmed at 4000 frames per second. Initially, the body orientation remained on the water surface with body angle (the body’s longitudinal axis (x’) relative to water surface), *θ*_*b*_ ≈ 0° (Fig. 1A, C-E). During this time, the whirligig moves parallel to the water surface at a speed of 45±17 cm/s (N=6). This is followed by the continuous change in *θ*_*b*_ after the head breaks the surface. At the end of diving, the body is reoriented by *θ*_*b*_ ≈ 90° with a considerable distance (1.3±0.3 body lengths) below the water surface (Fig. 1B, D-F).

**Fig. 1.**
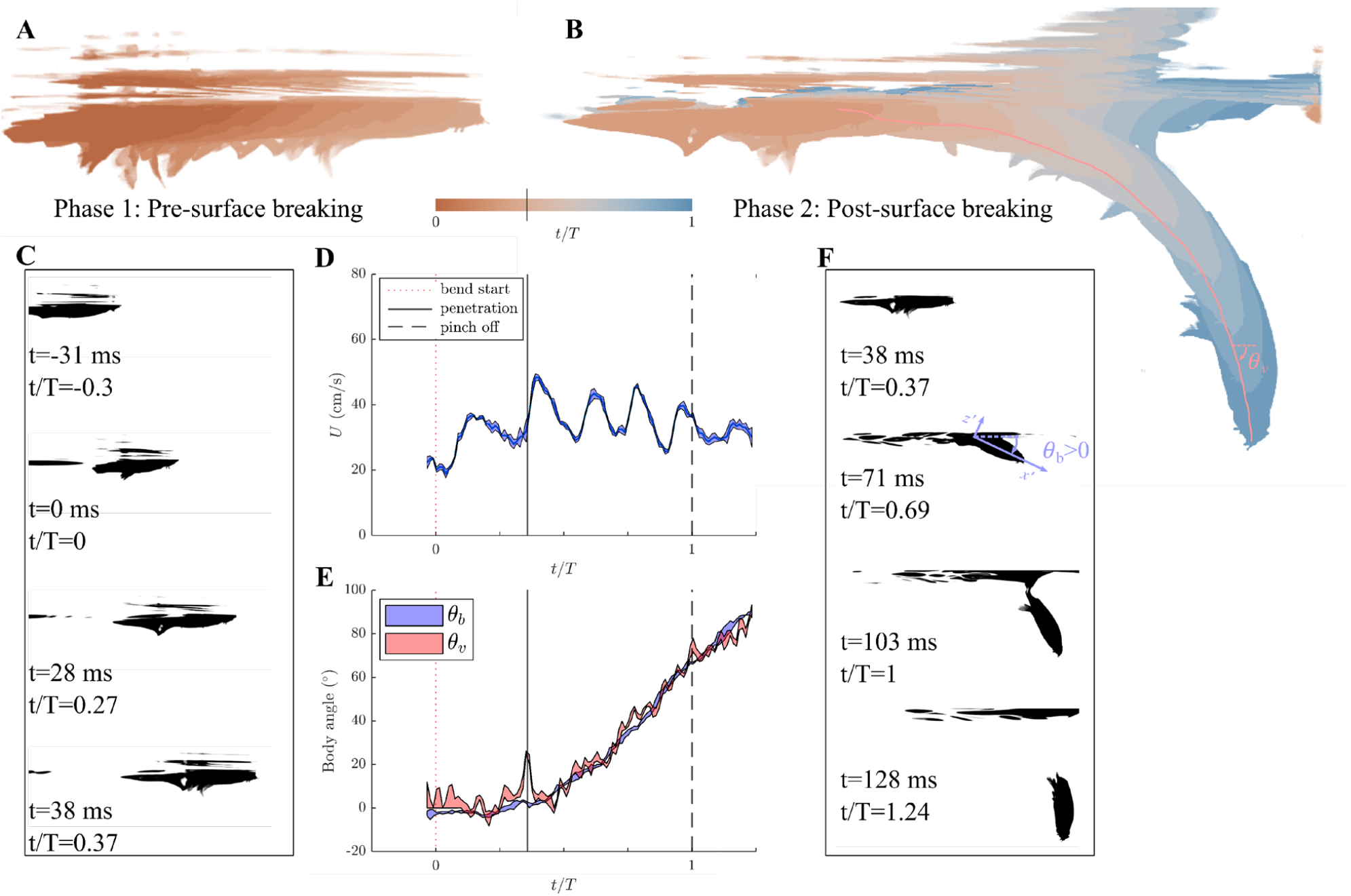
Kinematics of whirligig beetle in diving. The left side of the figure has information on phase 1 of diving (pre-surface breaking), and the right side of the figure has information on phase 2 of diving (post-surface breaking). (**A**) Diving Phase 1. (**B**) Diving Phase 2. (A-B) color corresponds to the color bar representing the time (*t*/*T*) normalized by the entire duration of the dive. (**C**) Phase 1 diving sequence. At t∼0 ms abdomen starts to bend. At t≈28 ms, the abdomen is nearly fully bent. At t≈38 ms, the head penetrates through the surface. (**D**) Time variation of the swimming speed, U. Phase 1 starts from the bend (red dotted line) to the surface penetration (black solid line). Phase 2 starts from the surface penetration to the air-water interface pinch off (black dashed line). (**E**) Time variation of the body angle, *θ*_*b*_ (defined in (F)) and the velocity direction angle, *θ*_*v*_ (defined in (B)). (**F**) Phase 2 diving sequence. At t∼103 ms, the air-water interface pinches off, and the beetle detaches from the surface. At t≈128 ms, the beetle’s body angle nears 90°.

We divided the diving process into two phases: pre-head penetration (phase 1; Fig. 1A, C-E) and post-head penetration (phase 2; Fig. 1B, D-F). In phase 1, although no appreciable body reorientation occurs, substantial torque is needed to break the water surface near the anterior end. In phase 2, the body begins to rapidly reorient as the dorsal surface progressively wets.

Prior to body re-orientation, we observed body morphing in the rear abdominal section (Fig.1; Movie S1). The last few segments (6-8th segments) of abdomen bent towards the ventral side (Fig. 1C t=28 ms). Such bending does not occur during their surface swimming (Movie S2). With the beginning of this notable conformation change as the starting point of diving, it took whirligigs 94±13 ms (N=6) to detach from the water surface (end of phase 2; Fig. 1F t=103 ms). The abdomen remains bent throughout the process, unbending near the end of phase 2.

### Surface resistance in Phase 1

To transition from phase 1 to phase 2, the beetle’s head needs to break through the water surface. When the head is pushed through the water, one can observe the water surface is effectively pinned around the outer hairs protruding from the labrum of the beetle (Fig. 2A-C, Movie S3), due to the small gap between the hairs (*21*). Once the head is pushed deep enough, the surface breaks and water rushes over the dorsal surface. With a contact angle of ≈90° (Fig. 2A), the upper surface provides little resistance to wetting. The submerging head with surface pinning generates an opposing vertical force, *F*_*head*_. To estimate this force, the hair and head of the whirligig beetle is approximated as a semi-circle with ultimate contact angle *ϕ* ≈ 90° (i.e., contact angle at which the surface breaks). Using the approximation from a previous study (*22*), the submergence depth of the head, *h*_*sub*_ at which the pinned contact angle reaches 90° is given by,

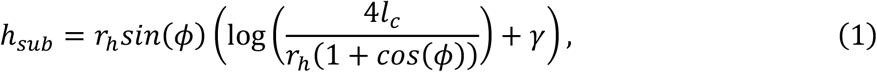

where *l*_*c*_ is the capillary length (∼2.7mm) and *r*_*h*_ is the radius of the head ∼ 1 mm, *γ* is the Euler constant, 0.577. The combined surface tension and buoyancy force would then be given by,

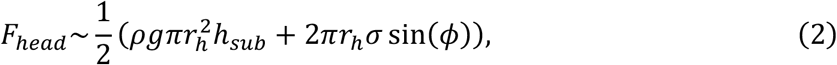

where *ρ* is the water density, g is the gravitational acceleration, *σ* is the surface tension. This results in *F*_*head*_ ≈ 0.25 *mN* resisting the body reorientation in phase 1.

**Fig. 2.**
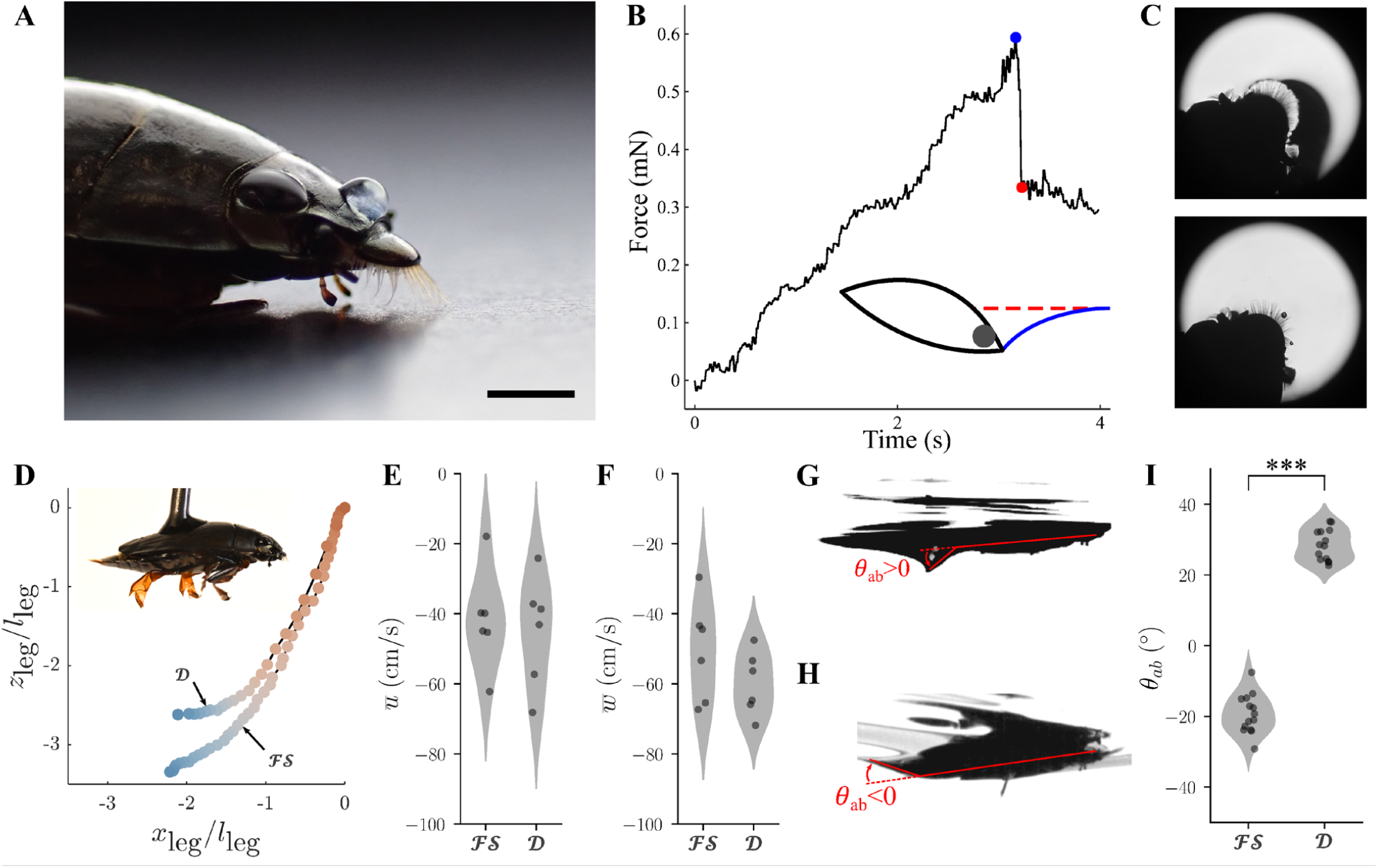
Resistance to surface breaking and comparative kinematics of interfacial swimming and diving. **(A)** Whirligig Beetle with labrum hair. The water droplet on its dorsal side shows near 90° contact angle. Scale bar = 1mm. **(B)** Water penetration force measurement. Schematic of the water surface shape before (blue solid line) and after (red dashed line) is shown in the left bottom corner. The sudden drop in the force between the blue circle and red circle is the relaxation of force resisted by head with hair. **C)** Labrum hair resisting surface penetration. Top image corresponds to pre-penetration indicated by the dark fringes around the hairs. Bottom image corresponds to post-penetration. (see also Movie S3) (**D**) Typical leg motion (normalized by leg length *l*_leg_) relative to body position in interfacial free swimming, **ℱ*S***, and diving, ***D***. From orange to blue, the color represents normalized time *t*/*T*_leg_, where *T*_leg_ is the duration of available measurements in leg stroke. Inset, a tethered live whirligig. (**E**) and (**F**) Distribution of time-averaged leg velocity 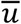 (*p*=0.73, *N*=6) and 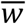 (*p*=0.21, *N*=6) relative to body in interfacial free swimming and diving, respectively. (**G**) and (**H**) Abdominal angle θ_ab_ (positive clockwise) relative to the longitudinal body axis in diving (***D***) and interfacial free swimming (**ℱ*S***), respectively. (**I**) Distribution of θ_ab_ (*p*∼10^−19^, *N*=14) in diving (***D***) and interfacial free swimming (**ℱ*S***).

To verify the estimated force and torque required to break the surface, a whirligig beetle tethered at 20° angle (tilted head down) was slowly pushed onto the water surface until the surface broke. The sudden drop in the vertical force was measured using a load cell (Futek LSB200 10g; See Supplementary Text for experimental details). When the bristle is removed, the measured force gradually changed indicating the role of the bristles in resiting the submergence (Fig. S1). The result shows that the head with hairs resist a force of 0.26±0.13 mN (N=5), similar to the theoretical approximation (Fig. 2B).

The theoretical and measured resistance values provide a conservative estimate of the force required to break the surface. When the beetle is diving, as the head pushes down on the water surface, flow would impinge on the depressed surface, which adds dynamic pressure to the static pressure considered. In turn, the surface would break more easily (i.e., the contact line would move over the hair at a lower static pressure). Thus, the static surface-breaking force establishes an upper bound for the required force.

Assuming the center of mass of the beetle is located at one-third of the body length from the head (a shift from the midpoint due to the large air-filled space under its elytra (*8*)), the pitching moment needs to exceed *τ*_*head*_*= l*_2_*F*_*head*_ ≈ 1 *mN*· *mm*, where *l*_2_ ≈ 4 *mN* is the distance between the head and the center of mass. This serves as the upper limit of the torque required. Next, the three potential sources of torque will be assessed, namely legs, anterior bending, and posterior bending.

### Leg as a potential source of torque in phase 1

For the leg to generate the torque for diving, its motion must be modified to alter the direction of the propulsive forces. During free-swimming, the leg motion generates torque that pitches the body upward. To generate opposite moment, the leg needs to generate the thrust above the center of gravity by moving the leg upwards, which is observed in other diving aquatic insects (*13*). However, Fig. 2D shows similar leg trajectories (*x*_leg_, *z*_leg_) in both interfacial swimming and diving. Moreover, the measured time-averaged leg velocities in both *x*- and *z*-directions 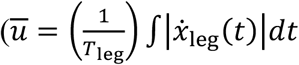 and 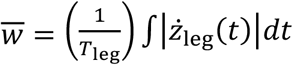, where *T*_leg_ is the duration of available measurement in leg stroke) shows similar leg velocities (Fig. 2E-F; Welch’s *t*(9.9)=0.36, *p*=0.73 for 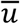 and *t*(8.4)=1.35, *p*=0.21 for 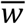, *N*=6). This further shows that no substantial modification of leg motion exists during whirligig’s diving. Thus, the leg is unlikely to provide the torque necessary for body rotation.

### Anterior bending as a potential source of torque in phase 1

For the dorsally dry anterior portion to generate the necessary torque, the head needs to protrude into the water and make a significant angle with respect to the rest of the body. Although the overall profile of the head is wedge shaped, and can provide pitching moment, the dry dorsal side in phase 1 experiences aerodynamic forces three orders of magnitude smaller than the hydrodynamic forces. As a result, only the conformation of the wetted ventral side can meaningfully contribute to the torque generation. During phase 1, the ventral side of the beetle’s anterior (*i*.*e*., head and prothorax) is flat (Fig. 2G, see Movie S1). While compared to the upward-curving conformation (similar to that of typical boat), this flat surface does prevent generation of pitch-up torque. Nonetheless, it cannot generate down-pitching torque necessary to initiate diving.

### Posterior bending as a potential source of torque in phase 1

Unlike its rigid dorsal side, the ventral side of whirligig beetle’s abdomen is flexible. Stimulation of the tethered whirligig shows that the whirligig can bend their abdomen up to 90° downward (Fig. S3). The abdominal shape in diving was significantly modified (Fig. 2G-H). The abdominal angle *θ*_*ab*_, the angle between the body axis and the abdomen (Fig. 2G-H), in diving and interfacial swimming are shown in Fig. 2I. A significant difference is observed in *θ*_*ab*_ (Welch’s *t*(24.7)=-25.30, *p*∼10^−19^), which even had a sign change between interfacial swimming (θ_ab_=-19±6°, *N*=14, upward bending) and diving (*θ*_*ab*_=28±4°, *N*=14, downward bending).

With the downward bending of the abdomen, the significant portion of the abdomen is exposed to the oncoming flow generating hydrodynamic forces. In turn, the torque about the center of mass would continuously generate the pitching down torque necessary to push the head down below the surface. Thus, the bent abdomen is most likely source of torque necessary for the transition from phase 1 to 2.

For a more precise comparison with the torque required for breaking the surface at the anterior end, a quantitative modelling of the torque generation by the posterior bending is required. In the following section, we model the hydrodynamic force and torque generated by the bent abdomen.

### Fluid force on the bent abdomen

At the air-water interface, the bent abdomen protruding into the flow experiences several different hydrodynamic forces, namely, viscous, steady inertial, unsteady inertial (added mass), surface tension, and buoyancy forces. The total force that the abdomen experiences, *F*_*abdm*_, can be expressed as,

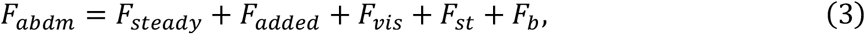

where *F*_*steady*_ is the steady inertial force, *F*_*added*_ is the unsteady inertial force, *F*_*vis*_ is the viscous force, *F*_*st*_ is the surface tension force, and *F*_*b*_ is the buoyancy force. To identify the dominant forces, the following dimensionless numbers comparing the forces are calculated,

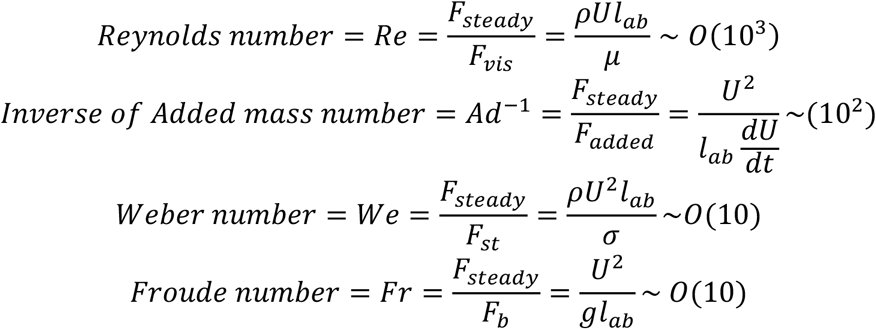

where *ρ* ≈ 1000*kg*/*m*^3^ is the density of water, *U* ≈ 0.45 *m*/ is the swimming speed, *L*_*ab*_ ≈ 3.3 *mm* is the typical length of the abdomen, µ ≈ 10^−3^kg/m/s is the dynamic viscosity of water, *dU*/*dt* ≈ 25*m*/*s*^2^ is the peak swimming acceleration, *σ*≈ 0.072 N/m is the surface tension, and *g* ≈ 9.8 *m*/*s*^2^ is the gravitational acceleration. These dimensionless numbers show that *F*_*steady*_ is the dominant contributor to *F*_*abdm*_, with its magnitude one to three orders of magnitude larger than the rest of the hydrodynamic forces. In the subsequent section, we model *F*_*steady*_ in detail.

### Steady Inertial Force and Torque

The most significant and continuous contribution to the torque originates from the steady inertial hydrodynamic force, *F*_*steady*_. To model *F*_*steady*_, we experimentally obtained the force coefficient, *C*_*F*_, at varying abdominal angles by combining flow visualization from the towing experiment and vortex panel method. The steady inertial force can be obtained by integrating the pressure around the wetted surface,

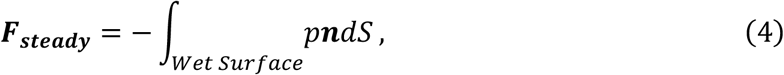

where p is the pressure, **n** is the outer normal unit vector.

A mechanical model with varying degrees of a sharply bent section was constructed and towed along the water surface to experimentally simulate phase 1 of the whirligig’s diving. The flow pattern was visualized with path lines constructed by combining consecutive particle images (Fig. 3A-C) (*23*). Note, because flow is steady, the path line (trajectory of the particle) is equivalent to the streamline (a line everywhere tangent to the local flow velocity).

**Fig. 3.**
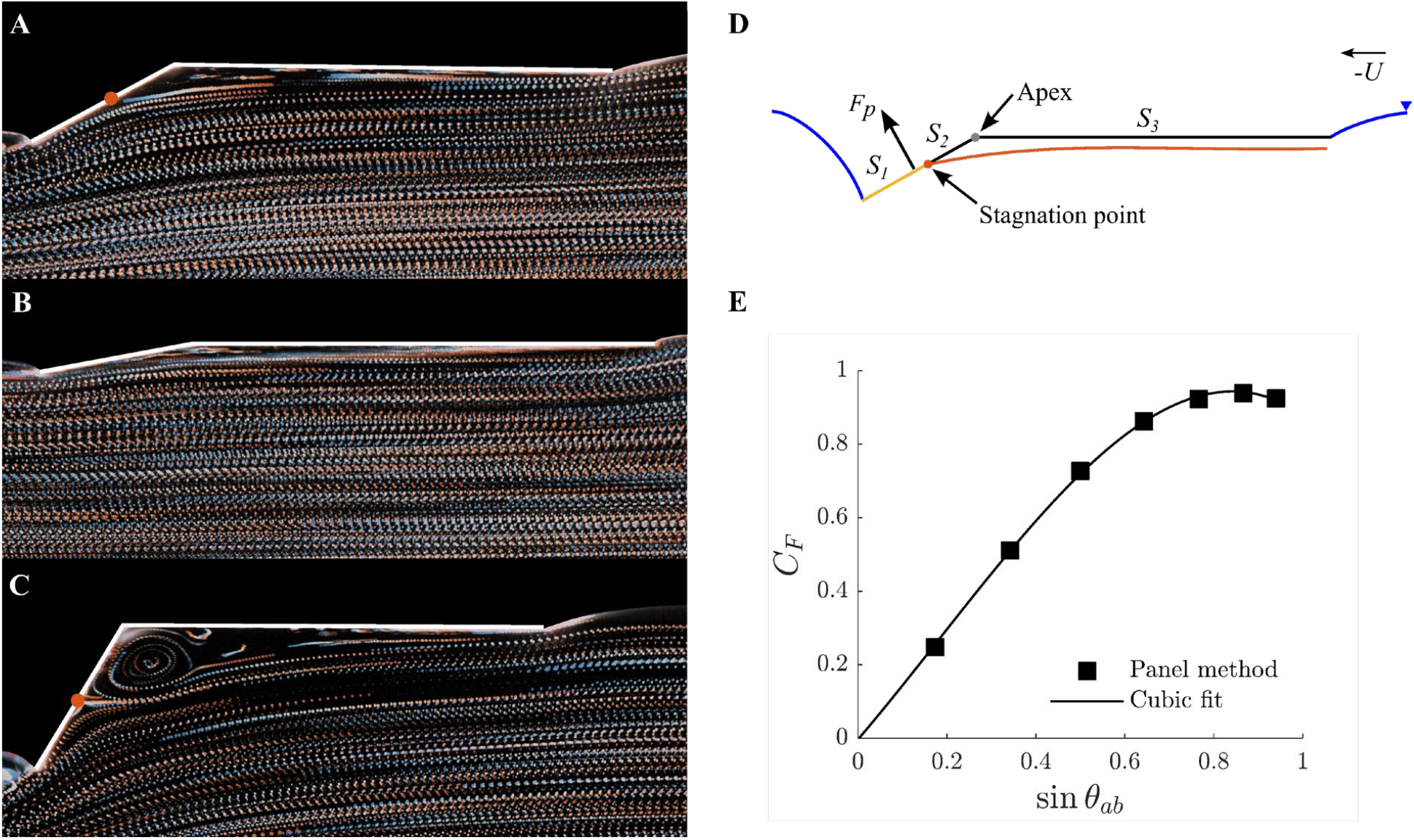
Flow visualization of towing experiments and results from numerical panel methods of the bent plate. (**A**—**C**) Path lines of flow under flat plate bent at (**A**) θ_ab_=30°, (**B**) θ_ab_=10°, and (**C**) θ_ab_=60°. Color code, sequential frame. Images were produced by combining 50 consecutive particle images. Red dots in (**A**) and (**C**), stagnation points. (**D**) Schematics of the mechanical model. The flat and bent sections are adjoined at the apex. The air-water interface (blue line) is pinned at the two ends of the plate. For θ_ab_<20°, the entire bent plate (black and yellow lines) was used as the wall geometry in the vortex panel method. For θ_ab_≥20°, the streamline (red line) terminating at the stagnation point (red dot) adjoined with the bent section (yellow line) downstream of the stagnation point is used as the wall geometry. The hydrodynamic pressure force ***F***_*p*_ acting normal to the bent section was assumed to be concentrated at its midpoint. Surface S_1_ is the wetted surface from the trailing edge to the stagnation point, S_2_ is from the stagnation point to the apex, and S_3_ is the flat surface. (**E**) Hydrodynamic pressure force coefficient *C*_*F*_ obtained from the vortex panel methods as a function of sin(θ_ab_). Black line, third-order polynomial fit.

At low abdominal angle (θ_ab_=10°, Fig. 3B), the streamlines remained parallel to the plate. As θ_ab_ increased (noticeably starting from θ_ab_=20°), the flow separated upstream of the apex (ridge of the fold; Fig. 3D) and reattaches onto the bent section. Fig. 3A shows the 30° case, which is close to the whirligig’s θ_ab_ (Fig. 2I). Here, the streamline terminating at the reattaching stagnation point (dividing streamline) clearly separates the inner region of stagnant flow from the outer region. As θ_ab_ further increased (θ_ab_=60°, Fig. 3C), flow separation became more pronounced, and the stagnation point moved further downstream. The recirculating flow inside the larger stagnant region is readily visible (Fig. 3C). These flow structures resemble classical flow over a forward-facing step, which has a high wall pressure inside the separation bubble (*24*).

The bent section is divided into two regions, *S*_*1*_ from the stagnation point to the trailing edge and *S*_*2*_ from the stagnation point to the bent point, i.e., apex (Fig. 3A). We implemented a potential flow model (Vortex Panel Method) (*25*) to calculate the pressure distribution along *S*_*1*_. For *S*_*2*_, we assumed the wall pressure as the uniform stagnation pressure ∼ρ*U*^2^/2, where *U* is the incident flow speed. For *θ*_*ab*_ < 20°, streamlines were assumed to follow the bent-plate geometry, therefore the potential flow model was used to calculate the pressure along the entire bent section.

The resulting pressure distribution was integrated along the bent section to find the net pressure force 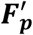 per unit width (see Supplementary Text for details), where 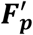 is defined as,

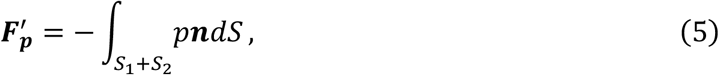

Non-dimensionalizing the 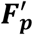 with inertial scaling gives *C*_*F*_,

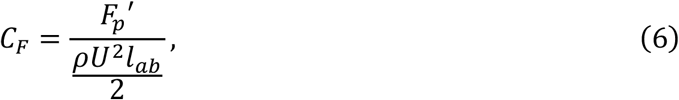

Because the projected area of the bent abdomen increases like sin(θ_ab_), we expect *C*_*F*_ to be a function of sin(θ_ab_) as well. Fig. 3E shows that *C*_*F*_ monotonically increases with sin(θ_ab_) up to about θ_ab_=60°. For the subsequent modelling effort, we used the third order polynomial fit to represent the relation *CC*_*F*_ = *f*(sin(*θ*_*ab*_)).

In addition, according to our flow visualizations in Fig. 3A-C, the stagnant region extends onto the flat section (unbent, parallel to horizontal water surface) spanning a length *l*_*s*_ when θ_ab_≥20°, where *l*_*s*_ is assumed equal to the length of the bent section inside the stagnant region (Fig. 4A; Supplementary Text). Therefore, the flat section inside the stagnant region experienced a hydrodynamic force (per unit width) 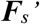, which is defined as

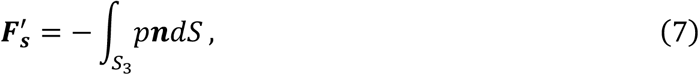

where S_3_ is the flat section. 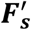 would have approximate magnitude of *ρU*^2^*l*_*s*_/2 according to Bernoulli’s principle. The total ***F***_***s****teady*_ is composed of both 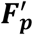 and 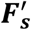, but the two forces act at different points.

Assuming that 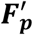 acts at the midpoint of the abdomen, the torque is given by 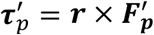, where r is the position vector from the center of mass to the abdomen midpoint (Fig. 4A). Further assuming that 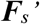 (zero for θ_ab_<20°) acts at the midpoint of this flat section (Fig. 4A), an additional upward force contributes to torque as 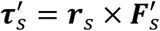, where ***r***_*s*_ is the lever arm of 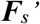 (Supplementary Text). Hence, the total body torque generated by the steady inertial force (per unit width) is represented as 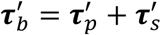.

**Fig. 4.**
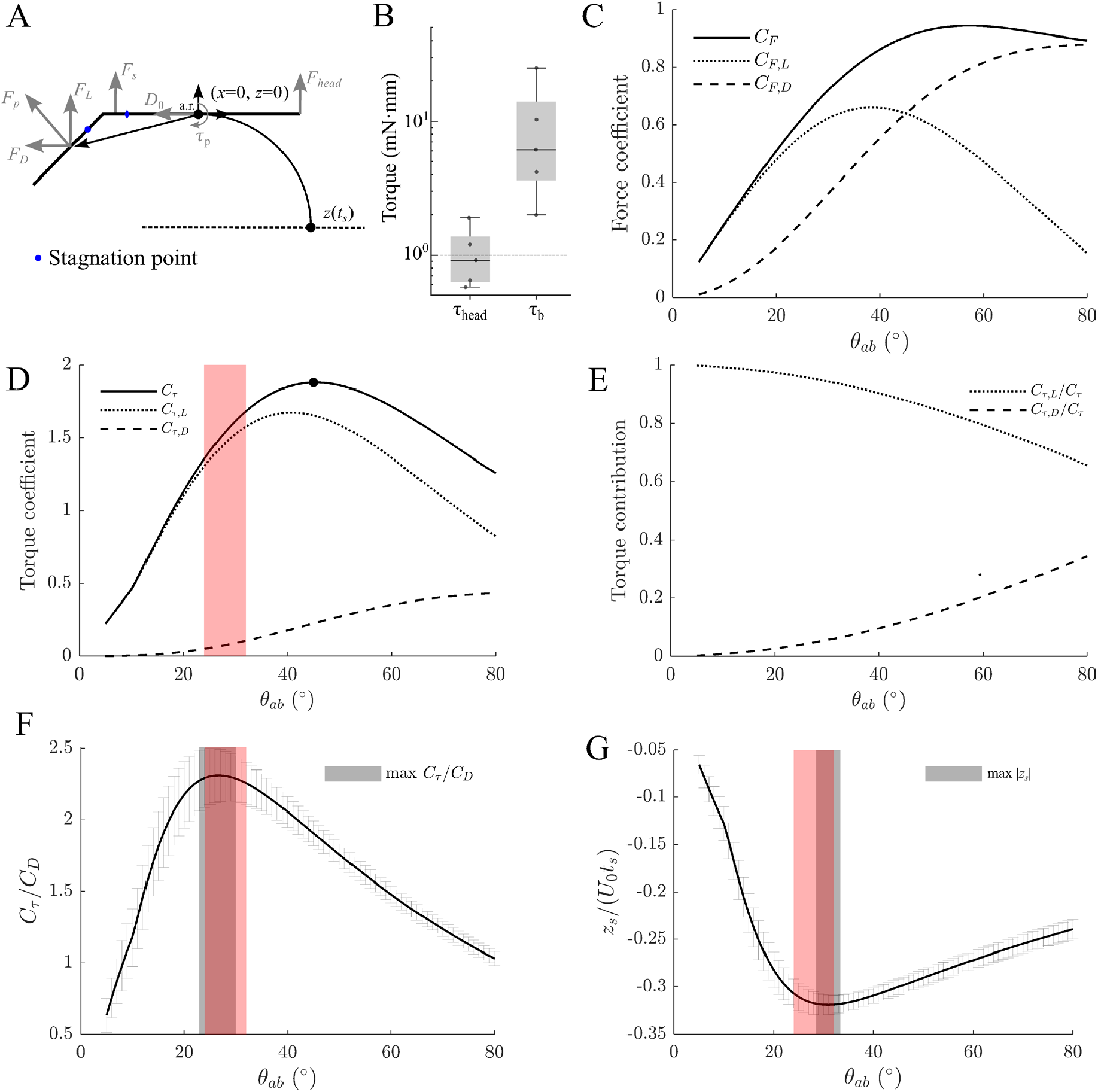
Schematics and results of the mechanistic model. (**A**) Schematics of forces and diving trajectory of the bent plate. The pressure force ***F***_*p*_, with components ***F***_*L*_ and ***F***_*D*_, acts at the midpoint of the bent section, forms a lever arm ***r*** from the axis of rotation, and produces torque ***τ***_*p*_ (positive clockwise). The flat section between the apex and the blue diamond is inside the stagnant region for θ_ab_≥20°. The distances from the apex to the blue diamond and the stagnation point (blue dot) are assumed equal. The baseline drag *D*_0_′ is assumed inline with the flat section. Dashed line, depth *z* reached after duration *t*_*s*_. (**B**) Comparison between *τ*_*head*_ and *τ*_*ss*_. The dotted line is the theoretical prediction of the *τ*_*head*_. The vertical axis is in log scale. (**C**) Force coefficients (*C*_*F*_, solid line; *C*_*F,L*_, dotted line; *C*_*F,D*_, dashed line) as functions of θ_ab_. (**D**) Coefficients of body torque (*C*_*τ*_, solid line) and its contributions from lift (*C*_*τ,L*_, dotted line) and drag (*C*_*τ,D*_, dashed line) as functions of θ_ab_. Black dot, maximum torque. (**E**) Relative torque contributions from lift (*C*_*τ,L*_/*C*_*τ*_, dotted line) and drag (*C*_*τ,D*_/*C*_*τ*_, dashed line) as functions of θ_ab_. (**F**) Torque-to-drag ratio (*C*_*τ*_*/C*_*D*_) as functions of θ_ab_. Solid line, average *C*_*τ*_*/C*_*D*_ across all values of *D*_0_′ and *β*. Gray shade, range of θ_ab_ for maximizing *C*_*τ*_*/C*_*D*_. (**G**) Depth *z*_*s*_ (normalized by *U*_0_*t*_*s*_) reached after duration *t*_*s*_ as functions of θ_ab_. Solid line, average *z*_*s*_ across all values of *D*_0_′and *β*. Gray shade, range of θ_ab_ for maximizing |*z*_*s*_|. Red shades in **E, F** and **G**, range of θ_ab_ from biological observations (28±4°, *N*=14). Error bars in **F** and **G**, one standard deviation.

Using the whirligig’s swimming speed and abdomen angle data with the mechanistic model, 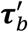 can be calculated. The results shows that the abdominal bending generates a torque 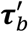 ranging from 2 to 25 mN·mm (9.5 ± 9.2 mN·mm, N = 5). The large variation is due to differences in swimming speeds and abdominal angles. This generated torque is well above the upper bound 1 mN·mm required to break through the water surface (Fig. 4B).

### Force and Torque decomposition

A deeper mechanistic insight can be gained through decomposing ***F***_*p*_*’* into two components in the direction perpendicular and parallel to the incident flow, namely lift ***F***_*L*_*’* and drag ***F***_*D*_*’* (Fig. 4A). While both components contribute to the torque, ***F***_*D*_*’* imposes a penalty by slowing down the beetle. With the same non-dimensionalization as in Eq. (4), the lift and drag coefficients (per unit width), *C*_*F,L*_=*C*_*F*_ cosθ_ab_ and *C*_*F,D*_=*C*_*F*_ sinθ_ab_, are shown in Fig. 4C. It shows that lift reaches maximum at θ_ab_≈39° and drag increases monotonically with θ_ab_ due to the increasing blockage of the incident flow

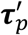 is decomposed in a similar way, 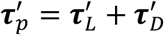 with 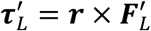 and 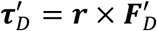, with the dimensionless torque coefficients defined as

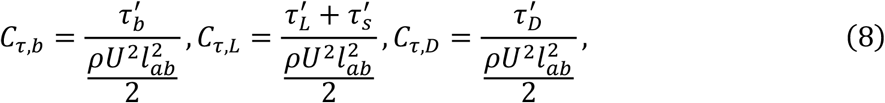

where *C*_*τ,b*_ represents the coefficient of total body torque (referred to as *C*_*τ*_ hereafter); and *C*_*τ,L*_ and *C*_*τ,D*_ represent the contributions from lift (on both the flat and the bent sections) and drag, respectively. The trends of the three torque coefficients with increasing θ_ab_ are shown in Fig. 4D. Both *C*_*τ,L*_ and *C*_*τ,D*_ followed similar trends as lift and drag on the abdomen (Fig. 4C)—*C*_*τ,L*_ peaked at θ_ab_≈41° and *C*_*τ,D*_ increased monotonically. With contributions from *C*_*τ,L*_ and *C*_*τ,D*_, the resulting body torque peaked at θ_ab_≈45°.

### Optimal Body Angle

A high torque enables faster and more reliable surface breaking in phase 1 and faster reorientation of the body in phase 2. This suggests that flexing the abdomen to θ_ab_≈45° is desirable for the whirligig beetle’s dive. However, the torque-maximizing θ_ab_ at 45° is 60% higher than the observed bending angle (∼28°, red shade in Fig. 4D; see also Fig. 2I). Thus, the maximizing torque does not appear to be the biological target for the whirligig beetle.

The observed abdominal angle could instead be due to the drag penalty associated with higher angles. As Fig. 4C shows, an increasing abdominal angle progressively increases the drag on the body. While the surface penetration and reorienting torque might occur faster, the drag leads to slower swimming speed, reducing the overall speed of their dive.

To elucidate the trade-off between torque generation and drag penalty, we further examined the relative contributions to body torque from lift and drag—*C*_*τ,L*_/*C*_*τ*_ and *C*_*τ,D*_/*C*_*τ*_—as shown in Fig. 4D. For a wide range of θ_ab_ (up to around the torque-maximizing optimum of 45°), lift dominates over drag in contributing to *C*_*τ*_, with *C*_*τ,L*_/*C*_*τ*_ an order of magnitude higher than *C*_*τ,D*_/*C*_*τ*_. Meanwhile, the relative contribution to *C*_*τ*_ from drag monotonically increases. The increasing drag penalty as θ_ab_ increases for higher torque represents a trade-off between torque generation and the associated drag.

The trade-off can be more clearly shown by computing an efficiency metric *C*_*τ*_/*C*_*D*_, the ratio of desired to undesired output. The *C*_*D*_ of the beetle can be modelled as

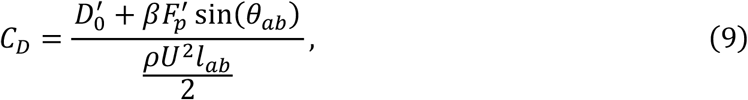

Where 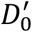 is a baseline drag per unit width, which accounts for the form drag on the body with zero-degree abdominal bending (2, 3), and β is a correction factor ranging from 0.8-1.2.

Fig. 4F shows the average *C*_*τ*_/*C*_*D*_ as a function of *θ*_*ab*_ with standard deviation across all 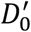 and *β. C*_*τ*_/*C*_*D*_ is maximized at *θ*_*ab*_ = 27° ± 2° (Fig. 4F, gray shade). At *θ*_*ab*_ smaller than the maximum, body drag is reduced but the low torque results in a smaller C_*τ*_/*C*_*D*_. In contrast, higher *θ*_*ab*_ increases torque while incurring increased drag penalty. Thus, *θ*_*ab*_ ≈ 27° represents optimally balanced angle, which closely overlaps with the *θ*_*ab*_ ≈ 28° observed in the whirligigs (Fig. 4F, red shade). Thus, their abdominal angle appears to target efficient torque generation rather than maximum torque generation.

The benefit of the optimal angle potentially extends to phase 2 of the diving. By rapidly re-orienting the body, while minimizing the speed loss due to the drag, the beetle may achieve faster diving. This will be explored in the next section.

### Role of posterior bending in Diving Phase 2

The phase 2 of the diving is considerably more complex as the dorsal surface begins to get wet. Unlike phase 1, the wedge-shaped anterior would progressively contribute to the pitching moment. While consideration of these additional hydrodynamic torque is necessary to fully describe the phase 2 diving kinematics, as a way to examine the role of the optimally bent abdomen in phase 2, we considered the equation of motion with abdomen generated torque as the only source of angular momentum. This simplified model can further elucidate the trade-off between the torque generation and drag penalty co-originating from the bent abdomen.

We modelled the diving trajectory for the duration between two leg strokes (*t*_*s*_ = 22±1 ms, *N*=6). In our model, body kinematics are prescribed by its position (*x,z*) and body angle θ_b_ (Fig. 4A). The diving starts at an initial speed *U*_*0*_ and body angle θ_b_=0°, and terminates at a depth of *z*_*s*_ after the duration *t*_*s*_. The body kinematics are governed by the conservation of linear and angular momentum (see Supplementary Text). We set the average initial speed observed (45 cm/s) as *U*_*0*_. Each combination of 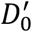 and *β* constructs an initial value problem, in which we solved for the terminal depth *z*_*s*_ (see Supplementary Text). Figure 4F shows *z*_*s*_ bounded by all combinations of 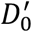 and correction factors for *F*_*p*_.

The results show that after the same duration *t*_*s*_, only small depth can be reached at small and large θ_ab_, both are associated with low *C*_*τ*_/*C*_*D*_ (Fig. 4F). We found that θ_ab_ for maximum *z*_*s*_ was 30°±2°, close to the biological abdominal angle and *C*_*τ*_/*C*_*D*_ maximizing angle. Thus, the near-optimal angle is not only efficient but also results in a faster overall dive.

### Application to Robotics

To demonstrate the feasibility of whirligig-inspired diving vehicle, a larger-scale whirligig-shaped model (8:1 scale) powered by two propellers (Fig. 5A) was constructed and tested (see Supplementary Text for more details on model construction and fabrication). The model’s outer contouring replicates a 3D model scan of a whirligig’s body. This model is equipped with an interchangeable control surface (i.e., flap) that mimics different abdominal angles (Fig. 5B-C, inset).

**Fig. 5.**
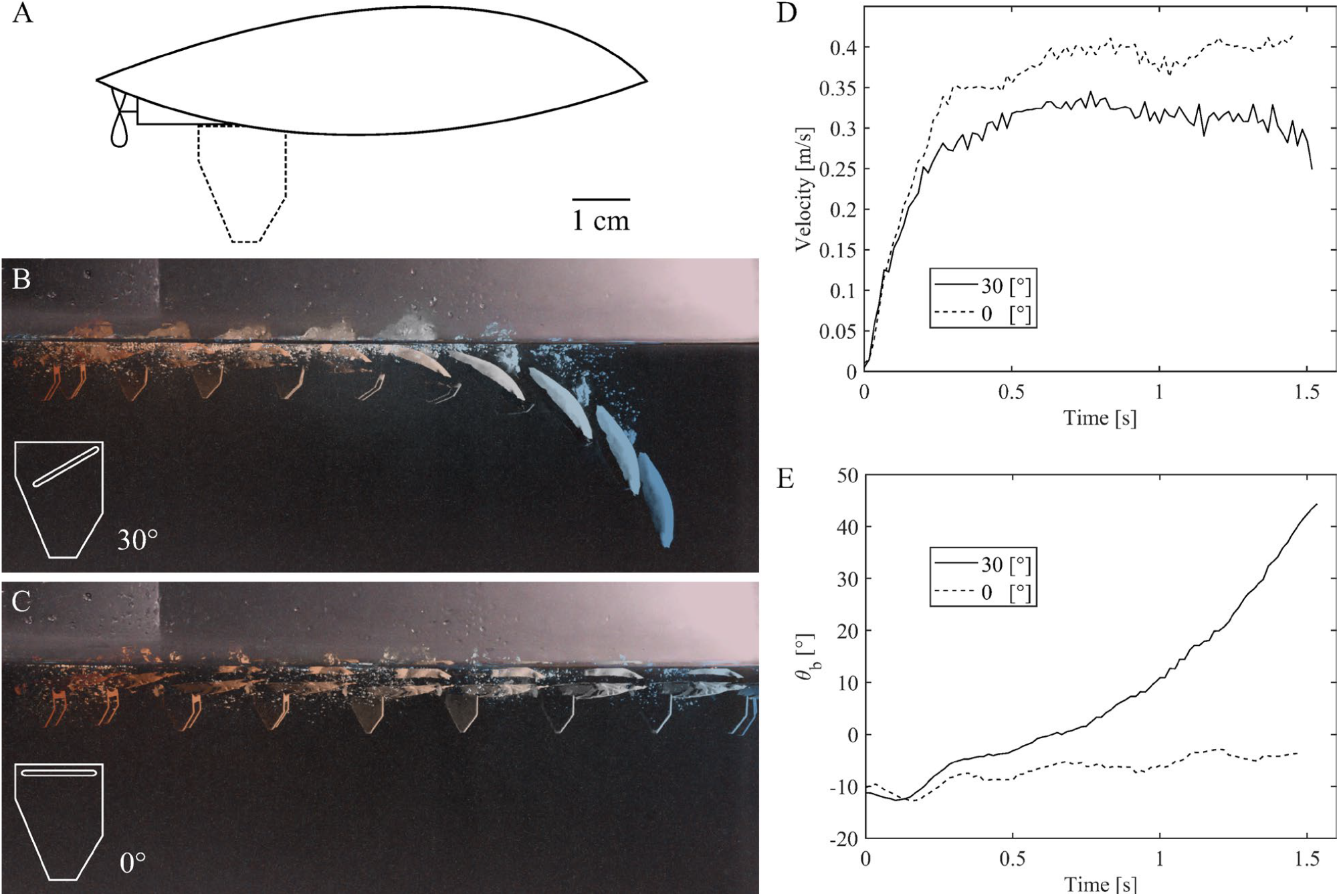
Mechanical model that mimics whirligig beetle diving. **(A)** Schematics of the 9.5-cm-long mechanical model, equipped with two propellers and an interchangeable control surface (dashed line). **(B-C)** Body trajectories with 30° and 0° flaps, respectively. Images (color-coded by time) were produced by combining 11 (for 30° flap) and 9 (for 0° flap) selected images from the Movie S4 and S5. Inset, cross-sectional schematics of the interchangeable control surface. **(D)** Temporal evolution of body speed of the model with 30° and 0° flaps. **(E)** Temporal evolution of body angle θ_b_ of the model with 30° and 0° flaps.

We filmed the side-view trajectories of the vehicle with 0° and 30° flaps (Fig. 5B-C, see Movies S4-5 for the full sequences). With the 30° flap, the vehicle underwent a downward-curved trajectory similar to that of the diving whirligig (cf. Figs. 1B and 2B). With the 0° flap, the vehicle’s motion remained on the water surface. Both had similar body speeds (Fig. 5D, small difference due to the additional drag incurred by the 30° flap), but the difference in body angle θ_b_ clearly shows that the flap angle can control the diving and non-diving states in a larger scale system (Fig. 5E). Due to its larger size, the model has minimal impediment from surface tension. Nevertheless, much like in the whirligig, the initial reorienting torque must come from the flap; otherwise, the vehicle with the 0° flap would have also dived.

## DISCUSSION

In this study, we examined the mechanism of the whirligig beetle’s rapid transition from free surface to underwater swimming. Our analysis of live whirligigs showed that whirligigs dive by morphing their body through abdominal bending. The bent abdomen results in increased hydrodynamic pressure that provides the pitching torque required to break through the water surface, which is resisted by the water surface effectively pinned to the hairs around their anterior edges.

Flow visualization of mechanical models along with pressure calculations allowed us to develop a torque model. The torque calculated from the model shows that the high swimming speed coupled with the bent abdomen provides sufficient torque.

The torque model further suggests that the beetle’s pitching torque is dominated by the lift force to avoid the drag penalty that slows them down. Similar to how Lift-to-Drag ratio is a measure of airfoil efficiency, maximization of *C*_*τ*_/*C*_*D*_ provides a way to optimally balance the trade-off between torque production and drag penalty. Thus, the beetle’s lift-based torque generation appears to be the most efficient form of diving.

### Other Sources of Torque during Diving Phase 2

The simplified mechanistic model of phase 2 diving showed that the abdominal angle employed by the beetle also results in faster diving. However, the phase 2 mechanistic model does not consider progressive dorsal wetting of the anterior portion. In addition to the continued torque by the flexed abdomen, the wedge-shaped head can also contribute to the reorientation process.

There are two notable changes that occur in phase 2. After the surface breaks, the labrum and hairs fold down, forming a sloped dorsal surface while the ventral side remains flat. Moreover, the top surface progressively wets as the beetle submerges. Note, the dorsal surface of the whirligig beetle is not strongly hydrophobic (the contact angle on the surface is approximately 90°); thus, a thin layer of air (plastron) that often forms on semi-aquatic insects does not form on the dorsal surface of the whirligig beetle.

As the beetle submerges with these changes, other parts of the body could start to contribute to the pitch-down moment hydrostatically and hydrodynamically. Hydrostatically, in the partially wetted configuration, the center of buoyancy and the center of mass are misaligned and produce hydrostatic torque until equilibrium is reached. Hydrodynamically, as the wedge-shaped head enters the water, the sloped dorsal surface experiences a lift force orthogonal to the direction of motion. Moreover, because the ventral surface is already submerged, the progressive wetting of the dorsal surface dynamically alters the added mass effect, similar to the case of the morphing body. As the beetle moves along its curved path, the advancing waterline over the sloped head displaces an increasing volume of fluid orthogonal to the beetle’s instantaneous trajectory. The rate of change in the fluid’s momentum exerts a corresponding reaction force in the opposite direction, pushing the anterior portion inward and further pitching the beetle down. Optimization of the head shape and entry kinematics is observed in sand burrowing lizards (*26, 27*), which “dives” into a granular material. A similar morphological optimization for water surface “burrowing” could have evolved for the whirligig as well.

### Re-visiting Past Observations and Hypotheses

The whirligigs’ diving behavior has been previously reported (*5*–*11*); however, these earlier studies did not identify the abdominal bending, which we showed as the key mechanism enabling the surface-breaking torque necessary to transition from interfacial swimming to diving.

In contrast to previous studies, the legs do not provide the primary pitching torque. Thrust vectoring by the legs that other studies observed is a consequence of the reoriented body, not the cause. In fact, close re-examining the diving sequences captured in one of the studies reveals clear abdominal bending preceding the dive (see reference (*5*) Xu et. al.’s figure 5), corroborating our findings.

Furthermore, the anterior bending suggested by Larsén (*3*), does not seem to directly impact surface penetration. The whirligig’s anterior portion is flat during phase 1, perhaps due to the strong resistance from the water line pinned around the hairs on the anterior margin. Without the protrusion into the water surface, the beetle is prevented from generating meaningful torque in the early phase of its diving. What Larsén might have observed is the continued reorientation in phase 2, which could visually appear as an anterior bending as the beetle progressively enters the water surface at a changing angle.

Larsén also provides a brief description of an ablation experiment noting the importance of the hindleg (*3*). The beetles with no hind-legs could not dive if their dorsal surface was dry, but were able to dive when the water was splashed over them. These results are consistent with the diving mechanism suggested in this work. The hind-legs are the main propulsor and are necessary for generating the fast speed that leads to sufficient hydrodynamic torque for breaking the surface. Swimming with mid-legs alone is too slow (achieving speed of ∼10 cm/s) and would fail to break the surface, preventing the whirligig beetles from diving. However, when water is splashed over the beetle, wetting the dorsal side of the beetle, it effectively removes the resistive barrier provided by the hair and surface tension; thus, allowing the beetle to dive even without the hindleg, albeit probably slowly.

### Ecological and Evolutionary Context and Robotics Translation

Although whirligig beetles are chemically well-defended (*28*), most assuring way to defend against predation is to flee. Between the two escape strategies, diving provides extended relief from predation compared to the rapid surface swimming. However, increased drag associated with their diving can temporarily increase the vulnerability of the beetle while they seek underwater refuge. The optimal abdominal angle that reduces this drag penalty allows them to transition to the safer underwater niche with reduced predation risk. Nevertheless, the cost of diving remains. While underwater, beetles cannot forage or seek mates, and are separated from the group (*29*–*31*). Thus, how they decide between fleeing on the water surface and diving remains to be studied.

For the semi-aquatic insects with wetted body, diving is a common niche-extension strategy. On the other hand, semi-aquatic insects that stand on the water surface with minimal wetting show counterpart escape adaptation: surface locomotion (*32*–*36*) and the interface-to-air transition (*36*–*38*). On land, whirligig’s diving escape is most similar to burrowing (*26, 27, 39*). Other similarly sized capable divers are diving beetles (*Dytiscidae*) and backswimmers (*Notonectidae*). They typically maintain a head-down posture at the interface, keeping the posterior tip of their abdomen in contact with the water surface for gas exchange (*19, 20, 40, 41*). Consequently, whereas whirligigs initiate dives from a flat, parallel posture on the water surface, their pre-tilted bodies allow them to readily direct thrust downward to move away from the surface. Furthermore, the relatively longer legs of *Dytiscidae* and water boatmen (*Corixidae*) allow for spatial adjustment of the thrust vector which can generate the forces and torque necessary for diving (*19, 20*). In contrast, the stout legs of whirligigs, which evolved for lift-based propulsion, likely constrained their ability to vector leg thrust, necessitating the evolution of a different diving strategy.

The downward abdominal bending aid in another niche transitions from land to air. When initiating their flight on land, whirligigs bend their abdomens to elevate their bodies (*3*). This provides ground clearance for wing deployment, compensating for their short hindlegs. Here, the beetle may fully bend their abdomens to maximize the body elevation. Thus, the simple bending mechanism facilitates two niche transitions: interface-to-underwater and land-to-air.

For the relatively large species of whirligig studied here (∼1 cm), the hydrodynamic pressure on the abdomen dominates the diving hydrodynamics. However, varying body size and speed observed across different species of whirligig beetles could change the relative importance of fluid forces. For example, the surface tension effect plays a relatively small role for the beetles tested in the current study as shown by the large Weber number (*We*∼*O*(10)). For a shorter body length, smaller *We* would indicate increasing importance of surface tension effect. The smallest whirligig beetle species with body length ∼3 mm (*42*), *Gyrinus cavatus*, operate at *We*∼*O*(1). The diving mechanism in this regime could experience both increased resistance to surface breaking and extra torque generation from the surface tension.

In addition to the downward-bent abdomen observed in the current study, other body morphing is observed in whirligigs. *Orectochilus* morph their abdomens laterally (in the plane perpendicular to *z*-axis, see Fig. 1A), acting as a ‘rudder’, which provides yawing torque for turning. Another species *Dineutus horni* unfolds their elytra for yawing motion (*43*). These examples highlight the broader potential of body-morphing control surfaces to provide complex maneuverability beyond pitching maneuvers. With the advances in soft and origami-inspired robotics (*44*–*48*), the more complex dynamic body morphing demonstrated in different whirligig species can be realized in robotic systems.

Robotic systems can further leverage unsteady effects during the body morphing such as added mass effects. The rate of change of angular momentum can be decomposed into 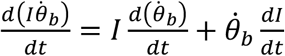, where *I* is the body moment of inertia. In a relatively slow bending of the whirligig’s abdomen, the d*I*/d*t* term is small. However, if the robotic system can rapidly morph its body (*49*), the d*I*/d*t* term can contribute significantly to the angular acceleration of the body. This kinematic strategy is observed in the aerial pitch control of jumping lizards (*50*), and has been implemented in robots with active tails that control the pitch dynamics (*51*), although the moment of inertia is provided mainly by the inertia of the tail rather than the surrounding fluid.

Diving maneuverability on the free surface provides whirligig beetles with an effective way to escape aerial predation by extending their niche to underwater. For the robotics community, this inspires a simple design solution for multimodal systems (*12, 48, 52*–*54*). Currently, some examples of robotic diving maneuvers include magnetic shape morphing (*48*), electrowetting to break the water surface (*52*), or asymmetric propulsion (*54, 55*). The one-degree-of-freedom bending of the whirligig’s abdomen provides an alternative using a control surface for a rapid pitch maneuver without modifying the propulsion mechanism. As demonstrated through the 8-to-1 mechanical model (Fig. 5), the diving mechanism shown here could be implemented in larger vehicles. We expect the whirligig-inspired solution to expand the operational domain of surface robots and surface vehicles to the underwater environment, paving the way for seamless multimodal systems.

## MATERIALS AND METHODS

### Statistical analysis

We performed Welch’s *t*-tests to compare the differences in leg velocities (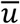 and 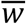) and abdominal angles θ_ab_ between interfacial free swimming and diving. The null hypotheses were 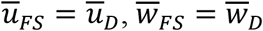, and θ_ab,*FS*_= θ_ab,*D*_, where the subscripts FS and D indicate interfacial free swimming and diving, respectively. The *t*-values for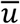, 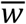 and θ_ab_ were *t*(9.9)=0.36, *t*(8.4)=1.35, and *t*(24.7)=-25.3, respectively. The *p*-values for 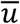 and 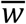 were *p*=0.73 and *p*=0.21, indicating that the leg velocities were similar in interfacial swimming and diving. However, the *p*-value for θ_ab_ was *p*∼10^−19^, suggesting that the abdominal angles were significantly different between interfacial swimming and diving.

## Acknowledgments

We thank Dr. Ronald Hoy, Dr. Mingming Wu and Dr. Christopher Dougherty for the insightful discussions about this study. We also thank Zhenghua Zhang for helping with prototyping the mechanical whirligig model. Y.S., S.J. and C.R. were supported by NSF-CMMI-2042740. Y.S. and C.R. were supported by NSF-CBET-2442036.

## Author contributions

Conceptualization: Y.S., C.R.

Methodology: Y.S., E.P., L.X., J.S., A.Q., C.R.

Investigation: Y.S., J.S., A.Q., C.R.

Visualization: Y.S., C.R.

Formal analysis: Y.S., S.J., C.R.

Supervision: C.R.

Writing—original draft: Y.S., C.R.

Writing—review & editing: Y.S., E.P., L.X., J.S., A.Q., S.J., C.R.

## Supplementary Materials

### The PDF file includes

Supplementary Text

Figs. S1 to S5

Table S1

Legends for Movies S1 to S5

### Other Supplementary Materials for this manuscript include the following

Movies S1 to S5

**·**

