## Supplementary Text for "Torque-optimized body morphing enables rapid surface to underwater diving in whirligig beetles"

### S1 Experimental details

#### S1.1 Diving kinematics

Whirligig beetles of genus *Dineutus discolor* were collected from a private pond in Ithaca, New York, USA. They were kept in a glass aquarium with pond water at room temperature. They were fed daily with dried crickets. Mass of live whirligigs after drying excess water were measured by a digital balance (ConicaCo SF-400C). The experiments were conducted in a 20cm × 20cm × 5cm glass tank elevated on aluminum beam structures and filled with tap water at 21°C. The side views of interfacial free swimming and diving were illuminated by an LED panel (Nanlite LumiPad 11). Diving was stimulated by intermittently placing hands between the free surface and an LED panel (Nanlite LumiPad 25) over the tank, such that the change in light intensity plays a similar role as the approaching predators (Fig. S2**A**).

Image sequences (side views) of diving behavior were collected by a high-speed camera (IDT Vision OS7) with a macro lens (Venus Optics Laowa 60 mm f/2.8). The camera was operating at 4000 frames per seconds with exposure time of 240 $\mu$s and a resolution of 1920 × 1080 pixels. The image recording was triggered digitally by motion capture feature in the image acquisition software (IDT Motion Studio). The pixel-to-length conversion was calibrated using a gridded glass slide placed in the focal plane.

##### S1.1.1 Kinematic data extraction

Image sequences only with horizontal swimming in the camera focal plane were analyzed ($N = 6$). The position of the anterior tip of whirligigs' body, $\mathbf{x}_b$, was hand tracked (Physlet Tracker 6.1.2) three times for each sequence. We measured the body angles ($\theta_b$) relative to horizontal water surface three times by hand tracking the contour of their body extracted using ImageJ 1.54d. To demonstrate the robustness of our measurement, we calculated the coefficient of variation, $CV = \sigma_{\theta_b}/\mu_{\theta_b}$, where $\sigma_{\theta_b}$ and $\mu_{\theta_b}$ are standard deviation and average of $\theta_b$ measured among the three repetitions. We found that $CV$ average over all measurements is 0.06, which indicates sufficiently small error. Because whirligigs carry air bubble under their elytra, the front is denser than the aft of the body. Therefore, we approximate the center of gravity (c.g.) was assumed to be located at one third of the body length.

Using the measured body angle $\theta_b$ averaged over the three repetitions, we calculated the position of c.g., denoted by $\mathbf{x}_b$, following the relation below:

$$\mathbf{x}_v = \mathbf{x}_g + \frac{l_b}{4}\left(\cos\theta_b\,\hat{\mathbf{e}}_x + \sin\theta_b\,\hat{\mathbf{e}}_z\right), \tag{S1.1}$$

where $\hat{\mathbf{e}}_x$ and $\hat{\mathbf{e}}_z$ are unit vectors in $(x, z)$ coordinates (see Fig. 1 in main text) and $l_b$ is the body length.

To increase the signal to noise ratio given our high sampling rate, We computed the body speed $\mathbf{u}_v$ at timestep $n$ using the centered difference scheme:

$$\mathbf{u}_v^n = \frac{\mathbf{x}_v^{n+5} - \mathbf{x}_v^{n-5}}{10\Delta t}. \tag{S1.2}$$

The trajectory angle was calculated as

$$\theta_v = \tan^{-1}\left(\frac{\mathbf{u}_v \cdot \hat{\mathbf{e}}_z}{\mathbf{u}_v \cdot \hat{\mathbf{e}}_x}\right). \tag{S1.3}$$

#### S1.2 Surface resistance experiment

Whirligig beetle was tethered at 20° angle (tilted head down) on a metal screw. The screw was then connected to a load cell (Futek LSB200 10g; Fig. S1A). The load cell was connected to single axis micro stage. The micro stage was slowly lowered pushing the beetle's head onto the water surface until the surface broke. The force was measured at 100 Hz sampling rate. A sudden drop in the force was measured to estimate the resistance force that originates from the water displaced by the surface pinning on the hairs (Fig. 2B). The same procedure was repeated using the beetle with its hair carefully removed using a sharp blade. The measured force did not show the sudden drop. Instead it gradual increased as the beetle submerges (Fig. S1B).

##### S1.3 Towing experiment

###### S1.3.1 Model design

The mechanical model was designed from a plate of width $w$ and total length $l_b$ with a section of length $l_{ab}$ bent at an angle $\theta_{ab}$ and a flat section of length $l_b - l_{ab}$ (Fig. S2**B**). The model was fabricated using a stereolithography (SLA) 3D printer (Formlabs Form 3+). To make sure that the contact line is pinned at the edge of the plate, the top surface of the model was coated with superhydrophobic material (NeverWet). The model was then placed approximately 0.4 mm below the undisturbed free surface to mimic the depressed meniscus of the free-floating beetles.

###### S1.3.2 Flow visualization

The model with various abdominal angles ($\theta_{ab} = 10°, 20°, ..., 70°$) towed at a constant speed of 23.6 cm/s in a 1 m × 0.3 m glass tank. The towing system (Fig. S2**C**) consisted of a linear rail (THK HSR-15C), a belt-driven carriage, and a stepper motor (Stepperonline Nema 23). The motor was supplied with 24V DC power and controlled by an Arduino Mega 2560 board using a digital motor driver (Stepperonline DM542T). The model was mounted on the rail using a 3D-printed mounting structure.

The bulk fluid was seeded with silver-coated glass particles (Potters Industries LLC, diameter 13 $\mu$m). The particles were illuminated by a 2-W 532 nm continuous laser (Ultralasers Inc), whose laser beam formed a laser sheet after passing through a combination of sheet optics (Thorlabs Inc). The laser sheet was ∼1 mm thick and was aligned with the mid-span of the model.

Image sequences of the flow motion were collected by a high-speed camera (Krontech Chronos 2.1) with a 100 mm f/2.8 lens (Venus Optics Laowa) operating at 1000 frames per second. The camera was fixed in space, meaning that the plate continuously moved past the camera view. To transform to the frame of reference of the plate, images were shifted at the towing speed using sub-pixel linear interpolation (MATLAB R2024b). Path lines of the flow were obtained by combining 50 consecutive images. To indicate flow direction, particle images were color-coded temporally. Further details can be found in our previous work (*16*).

#### S2 Calculations of wall pressure

We implemented the two-dimensional vortex panel method for solving lifting airfoil problems. In this method, the geometry combining the dividing streamline and the bent section (Fig. 2**D** in the main text) is discretized into $N_p$ panels. Each panel represents a vortex sheet with uniform strength (circulation) $\Gamma$, such that each discrete vortex induces a flow velocity at all other panels, including its own, according to the Biot-Savart law (*13*). To ensure that Kutta condition is satisfied at the posterior end of the plate, circulation and velocity induced by all vortices are stored at the quarter-chord and midpoint of each panel.

Denote velocity stored at each panel as $\mathbf{u}_i$ and circulation as $\Gamma_j$. The velocity $\mathbf{u}_i = u_i\hat{\mathbf{e}}_x + w_i\hat{\mathbf{e}}_z$ at each panel composes of velocities induced by the vortices at all panels is given by (*18*):

$$\begin{aligned} u_i &= \sum_j^{N_p} \frac{\Gamma_j}{2\pi}\frac{(z_i - z_j)}{r_{ij}^2}, \\ w_i &= \sum_j^{N_p} \frac{-\Gamma_j}{2\pi}\frac{(x_i - x_j)}{r_{ij}^2}, \end{aligned} \tag{S2.1}$$

where $j$ represents the velocity induced by the vortex located at the $j-$th panel, and $r_{ij}^2 = (x_i - x_j)^2 + (z_i - z_j)^2$ is the distance between the two panels. At the panels, no-penetration boundary condition is satisfied:

$$(\mathbf{u} + \mathbf{u}_\infty) \cdot \hat{\mathbf{n}} = 0, \tag{S2.2}$$

where $\mathbf{u}_\infty = U_\infty \hat{\mathbf{e}}_x$ and $\hat{\mathbf{n}}$ is the outward normal at the plate. Eq. S2.2 therefore turns into $N_p$ algebraic equations numerically solved for $\Gamma_j$ using MATLAB R2024b.

With the computed vortex strength, the tangential component of the induced velocity, $\mathbf{u}_t = (\mathbf{u} + \mathbf{u}_\infty) \cdot \hat{\mathbf{t}}$, where $\hat{\mathbf{t}}$ is the unit tangential of the panel, can be obtained. Here $\mathbf{u}_t$ represents the principal-value tangential velocity on the vortex sheet (*18*) and is therefore the average between flow velocities slightly above and below the vortex sheet, denoted as $\mathbf{u}^+$ and $\mathbf{u}^-$, respectively, which are in equal magnitude and opposite directions. With the computed vortex strength $\Gamma$, which represents the velocity jump across the vortex sheet, the flow velocity slightly below the vortex sheet is computed as

$$\mathbf{u}^- = \mathbf{u}_t - \frac{\Gamma}{2\Delta s}, \tag{S2.3}$$

where $\Delta s$ is the length of the panel. Using Bernoulli equation for small-disturbance flow over an airfoil, the pressure along the lower surface is computed as:

$$p^- - p_\infty = -\rho U_\infty |\mathbf{u}^-|. \tag{S2.4}$$

The magnitude of the hydrodynamic pressure force $\boldsymbol{F}_p$ acting on the bent section of the plate was therefore

$$F_p = \begin{cases} \sum\limits_{i \in C_1 + C_2} p_i^- \Delta s_i, & \theta_{ab} < 20°, \\ \sum\limits_{i \in C_1} \frac{1}{2}\rho U_\infty^2 \Delta s_i + \sum\limits_{i \in C_2} p_i^- \Delta s_i, & \theta_{ab} \geq 20°, \end{cases} \tag{S2.5}$$

where $C_1$ and $C_2$ are the section from the apex to the stagnation point and the remaining section it, respectively (see Fig. 2**D** in main text).

The plate geometry is treated as follows. For $\theta_{ab} < 20°$, we used the entire plate as plate geometry. For $\theta_{ab} \geq 20°$, we manually extracted the shape of the streamline terminating at the stagnation point (using ImageJ 1.54d) adjoined with the remaining bent section of the plate. To smooth the streamlines, their extracted coordinates were fitted with third-order polynomials using `polyfit` in MATLAB R2024b.

The hydrodynamic pressure force coefficient, $C_F$, is obtained using Eq. 1 in the main text. We fitted a third-order polynomial to $C_F$ (see Fig. 2**E** in the main text):

$$C_F = A_0 + A_1 \sin\theta_{ab} + A_2 \sin^2\theta_{ab} + A_3 \sin^3\theta_{ab} \tag{S2.6}$$

and $A_0 = 0$, $A_1 = 1.42$, $A_2 = 0.65$ and $A_3 = -1.18$ were obtained from fitting.

For $\theta_{ab} \geq 20°$, the length of the flat section inside the stagnant region was assumed equivalent to the distance between the apex and the stagnation point (see Fig. 3A in the main text), denoted as $l_s$, which was obtained from the endpoints of the extracted streamlines. Similar to $C_F$, we fitted $l_s$ as a function of $\sin\theta_{ab}$ using a third-order polynomial:

$$\frac{l_s}{l_{ab}} = A_{s0} + A_{s1} \sin\theta_{ab} + A_{s2} \sin^2\theta_{ab} + A_{s3} \sin^3\theta_{ab}, \tag{S2.7}$$

where $A_{s0} = -0.21$, $A_{s1} = 1.02$, $A_{s2} = 1.01$ and $A_{s3} = -1.22$.

The pressure along the bent section between the apex and the stagnation point and the flat section spanning length $l_s$ were approximated as the stagnation pressure $\rho U_\infty^2/2$.

#### S3 Mechanistic model

In this model, we consider the dynamic forces on the body during diving (Fig. 3A in the main text). We use the same geometry as used in the previous section. However, the forces and torques in this section are three dimensional and are calculated based on morphological measurements of whirligig beetle. We denote body length by $l_b$, body width by $l_w$, body height as $l_h$, and body mass as $M$, whose corresponding values are shown in Table S1.

The body experiences a baseline drag $D_0$ at the center of rotation and hydrodynamic pressure force $F_p$ at the midpoint of the abdomen. The baseline drag is defined as

$$D_0 = C_{D0} \frac{1}{2} \rho U^2 S_f, \tag{S3.1}$$

where $C_{D0}$ is the baseline drag coefficient and $S_f = \pi l_w l_h / 8$ is the half front area of the whirligig (assuming elliptical cross section). Given $Re \sim O(10^3)$, we varied $C_{D0}$ between 0.4 and 0.6.

The hydrodynamic pressure force $F_p$ is defined as

$$F_p = C_F \frac{1}{2} \rho U^2 S_{ab}, \tag{S3.2}$$

where $S_{ab}$ is the abdomen area:

$$\begin{aligned} S_{ab} &= \int_{l_b/2-l_{ab}}^{l_{ab}/2} l_w \sqrt{1-\left(\frac{2x}{l_b}\right)^2}\, dx \\ &= \frac{l_w l_b}{4}\left\{\frac{\pi}{2} - \left[\left(1-\frac{2l_{ab}}{l_b}\right)\sqrt{1-\left(1-\frac{2l_{ab}}{l_b}\right)^2} + \arcsin\left(1-\frac{2l_{ab}}{l_b}\right)\right]\right\}. \end{aligned} \tag{S3.3}$$

The total drag on the body is therefore

$$D = D_0 + \beta F_p \sin\theta_{ab}, \tag{S3.4}$$

where $\beta$ is the correction factor varied between 0.8 and 1.2.

Assuming the axis of rotation is at the midpoint of the flat section of the body, the lever arm $\mathbf{r}$ of $F_p$ in $x-$ and $z-$ directions are decomposed as:

$$\begin{aligned} r_x &= \frac{l_b - l_{ab}}{2} + \frac{l_{ab}}{2}\cos\theta_{ab}, \\ r_z &= \frac{l_{ab}}{2}\sin\theta_{ab}. \end{aligned} \tag{S3.5}$$

The torque contributed from the hydrodynamic pressure force on the abdomen is

$$\tau_p = \beta F_p (r_x \cos\theta_{ab} + r_z \sin\theta_{ab}). \tag{S3.6}$$

For $\theta_{ab} \geq 20°$, the hydrodynamic pressure force $F_s$ on the flat section spanning length $l_s$ is defined as:

$$F_s = \frac{1}{2}\rho U^2 S_{\text{flat}}, \tag{S3.7}$$

where $S_{\text{stag}}$ is the area of the flat section spanning length $l_s$, assuming the same planform of the body as in obtaining $S_{ab}$. Therefore, $S_{\text{flat}}$ is found by evaluting the integral in Eq. S3.3 from $l_b/2 - l_{ab} - l_s$ to $l_b/2 - l_{ab}$:

$$\begin{aligned} S_{\text{flat}} = \frac{l_w l_b}{4} \Bigg\{ &\left(1-\frac{2l_{ab}}{l_b}\right)\sqrt{1-\left(1-\frac{2l_{ab}}{l_b}\right)^2} + \arcsin\left(1-\frac{2l_{ab}}{l_b}\right) \\ &- \left[1-\frac{2(l_{ab}+l_s)}{l_b}\right]\sqrt{1-\left[1-\frac{2(l_{ab}+l_s)}{l_b}\right]^2} - \arcsin\left[1-\frac{2(l_{ab}+l_s)}{l_b}\right]\Bigg\}. \end{aligned} \tag{S3.8}$$

Assuming that $F_s$ acts perpendicular to the midpoint of $l_s$, the lever arm formed by $F_s$ is:

$$r_s = \frac{1}{2}(l_b - l_{ab} - l_s). \tag{S3.9}$$

The torque contributed from the hydrodynamic pressure force on the flat section is

$$\tau_s = r_s F_s. \tag{S3.10}$$

The total torque on the body is therefore

$$\tau_b = \tau_p + \tau_s. \tag{S3.11}$$

Note, Eqs. S3.5 and S3.9 also hold for the two-dimensional approximation (for force per unit width).

The moment of inertia of the body varies with the abdominal angle $\theta_{ab}$ and is in general defined as:

$$I = \int r^2\, dm, \tag{S3.12}$$

where $r$ is the distance from the axis of rotation to $dm$. Assuming uniform mass distribution, we have $dm = (M/l_b)ds$ and the integral in (S3.12) becomes

$$I = \frac{M}{l_b}\int r^2(s)\, ds. \tag{S3.13}$$

The moment of inertia is calculated by adding up the integrals evaluated for the flat and the bent sections. In the horizontal section:

$$r^2 = s^2, \text{ where } r \in \left[-\frac{l_b - l_{ab}}{2}, \frac{l_b - l_{ab}}{2}\right]. \tag{S3.14}$$

In the bent section:

$$r^2 = \left[\frac{l_b - l_{ab}}{2} + s \cdot \cos\theta_{ab}\right]^2 + (s \cdot \sin\theta_{ab})^2, \text{ where } s \in [0, l_{ab}]. \tag{S3.15}$$

By evaluating (S3.12) with integration parameters in (S3.14) and (S3.15), the moment of inertia for rotation about the axis of rotation is

$$I = \frac{M}{l_b}\left[\frac{2}{3}\left(\frac{l_b - l_{ab}}{2}\right)^3 + \frac{l_{ab}(l_b - l_{ab})}{2}\left(\frac{l_b - l_{ab}}{2} + l_{ab}\cos\theta_{ab}\right) + \frac{1}{3}l_{ab}^3\right]. \tag{S3.16}$$

The equations of motion are given by

$$M\frac{dU}{dt} = -(D_0 + \beta F_p \sin\theta_{ab}), \tag{S3.17a}$$

$$I\frac{d\dot{\theta}_b}{dt} = \tau_b. \tag{S3.17b}$$

where $D_0$, $F_p$, $\tau_b$ are all functions of time. Eqs. S3.17a and S3.17b were integrated with time from $t = 0$ to $t = t_s$ using first-order Euler scheme. Here $t_s = 22$ ms is the typical stroke-to-stroke duration measured from the image sequences. Next, we applied the following kinematic relations:

$$\frac{dx}{dt} = U\cos(-\theta_b), \tag{S3.18a}$$

$$\frac{dz}{dt} = U\sin(-\theta_b), \tag{S3.18b}$$

$$\frac{d\theta_b}{dt} = \dot{\theta}_b. \tag{S3.18c}$$

The depth $z_s$ was obtained by integrating Eqs. S3.18a, S3.18b and S3.18c using the same numerical scheme.

##### S3.1 Forces and torques per unit width

The forces and torques per unit width examined in the "Results" section of the main text were obtained by normalizing with a characteristic width of the abdomen $w_{ab}$ as:

$$(\cdot)' = \frac{(\cdot)}{w_{ab}}, \tag{S3.19}$$

where $w_{ab} = S_{ab}/l_{ab}$ and $(\cdot)$ represents the fores ($F_p$ and $D$) and torques ($\tau_p$, $\tau_s$ and $\tau_b$) computed in the previous section, and $(\cdot)'$ represents the corresponding per-unit-width values.

##### S3.2 Parameter space

The morphological measurements of the whirligig and the ranges of parameters used in our dynamic model are given in Table S1.

#### S4 Mechanical whirligig model in Fig. 5

An up-scaled mechanical model of the whirligig was constructed. The shape of the model was obtained from nano-computed tomography (nano-CT) imaging of a deceased whirligig. The specimen was placed in Tygon tube and imaged at a resolution of 6.96 $\mu$m using a nano-CT scanner (Zeiss-Xradia Versa 520 X-ray microscope). The CT images were first converted into surface meshes, which were then mapped into a solid body containing a top and a bottom shell (see Fig. S4) in Solidworks 2025. The model was 3D printed using an SLA 3D printer (Formlabs Form 4). The propellers and micro controller of the model were obtained

| Category | Description | Symbol | Range or Value |
|---|---|---|---|
| Morphological | Body length (mm) | $l_b$ | $11.85 \pm 0.50$ |
| | Body width (mm) | $l_w$ | $6.56 \pm 0.32$ |
| | Body height (mm) | $l_h$ | 3.73 |
| | Abdomen length (mm) | $l_{ab}$ | $3.28 \pm 0.14$ |
| | Body mass (mg) | $M$ | 55 |
| Mechanistic model | Baseline drag coefficient (frontal area) | $C_{D,0}$ | $0.4 - 0.6$ |
| | Correction factor for $F_p$ | $\beta$ | $0.8 - 1.2$ |

**Table S1.** Morphological measurements and ranges of parameters used in the mechanistic model.

from an RC control boat (PowerTRC RH711). The model was equipped with 3D printed, interchangeable control surfaces (flaps with angles of 0° and 30°, see Fig. S5).

Side-view trajectories of the model moving in a glass aquarium were filmed by a mirrorless camera (Nikon Z5) with a macro lens (Venus Optics Laowa 160 mm f/1.8) at 60 frames per second. Features on the model were tracked to extract its trajectory, body speed and body angle $\theta_b$, following the same procedures in Section S1.

### Supplementary Figures

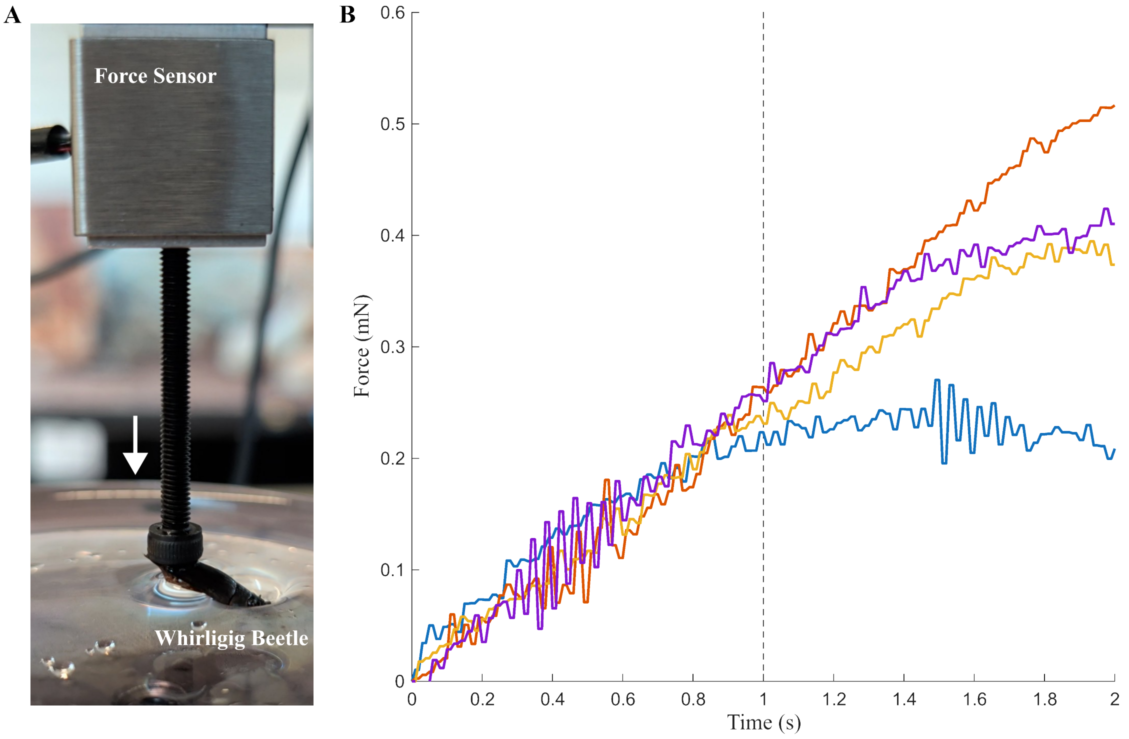


**Fig. S1.** Experimental setup for surface resistance force measure. (A) Experimental setup with whirligig's head pushing on the water surface. (B) Force data of the beetle lowered with hair removed. Water line started wetting the top surface around time = 1s (vertical line). Four different measurement show no sudden drop in forces at time = 1s.

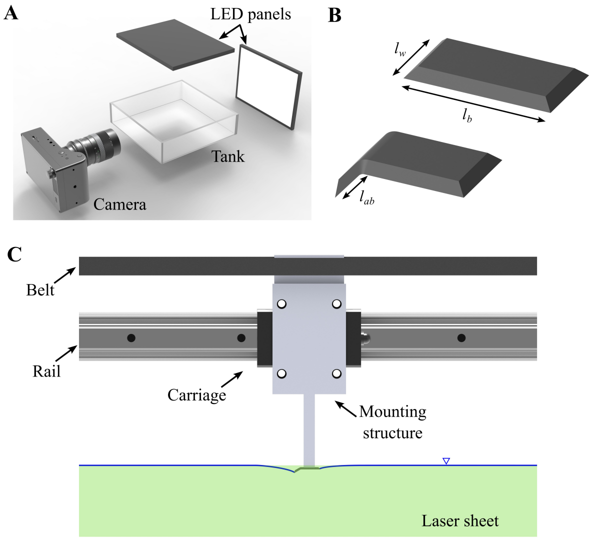


**Fig. S2.** Schematic illustrations of experimental setups. (A) Setup for diving experiments. Whirligig beetles in interfacial free swimming are filmed in side view. The side view and water surface are illuminated by LED panels. (B) Design of mechanical model for flow visualization. Comparison of bent and unbent plates is shown. The width and total length of the plate is $w$ and $l_b$, respectively. The length of the bent section is $l_{ab}$. (C) Setup of towing system. The bent plate is mounted to the carriage through a 3D-printed mounting structure. The carriage is moved along a rail through a belt driven by a stepper motor.

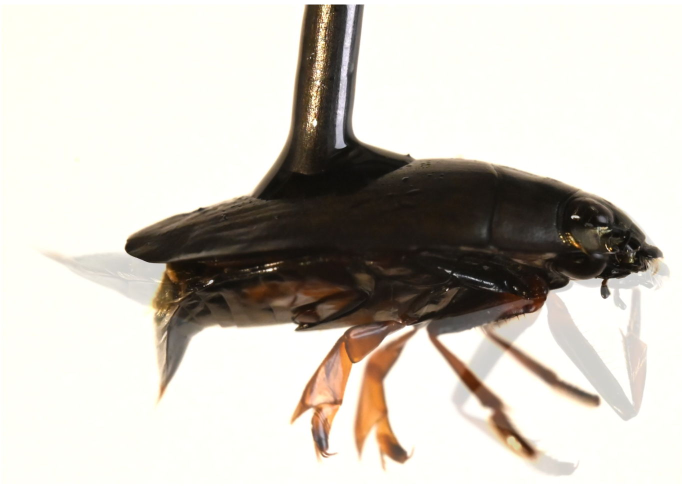

**Fig. S3.** A tethered whirligig beetle. The whirligig's abdomen is bent at $\sim 90°$ when stimulated. The image is shadowed with the unstimulated whirligig.

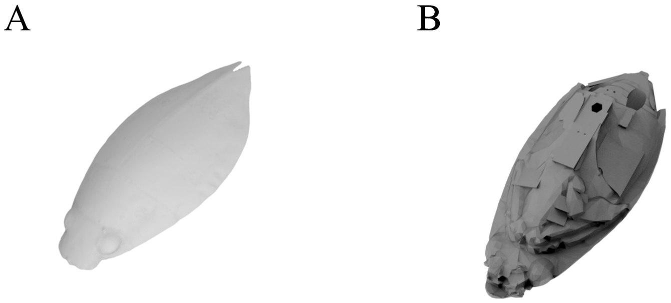

**Fig. S4.** CAD model of the mechanical whirligig consisting of two a top and a bottom shell. (A) Top shell. (B) Bottom shell.

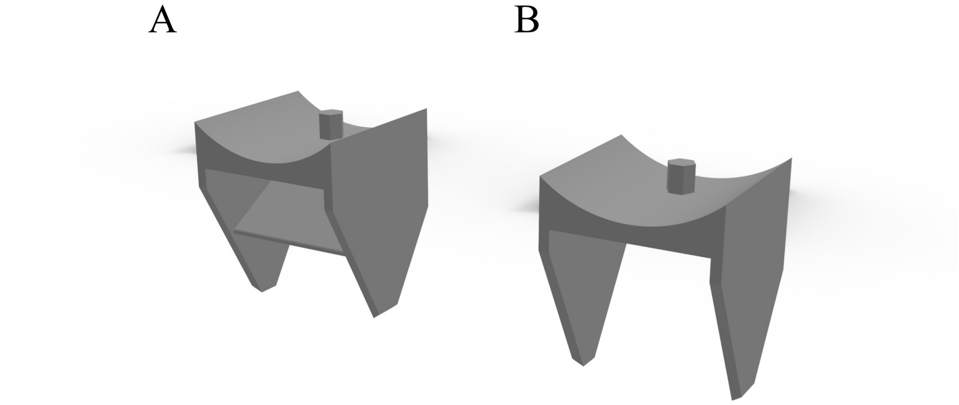

**Fig. S5.** CAD model of the flap of the mechanical whirligig. (A) 30°. (B) 0°.

#### Supplementary Movies

##### Movie S1.

Supplementary movie of whirligig beetle diving from free-surface swimming (playback speed, 40 fps, 100x slower than real time).

##### Movie S2.

Supplementary movie of whirligig beetle upward-pitching during leg power stroke (playback speed, 40 fps, 100x slower than real time).

##### Movie S3.

Supplementary movie of whirligig beetle's head pushing down onto the water surface. (playback speed, 60 fps, real-time speed)

##### Movie S4.

Supplementary movie of mechanical surface vehicle with flap oriented at 0 degrees (playback speed, 60 fps, real-time speed).

##### Movie S5.

Supplementary movie of mechanical surface vehicle with flap oriented at 30 degrees (playback speed, 60 fps, real-time speed).
